# Novel humanized anti-Nav1.7 antibodies with long-lasting, side-effect-free analgesic effects

**DOI:** 10.1101/2025.08.22.671720

**Authors:** Sosuke Yoneda, Daisuke Uta, Kana Yasufuku, Takuya Yamane, Saho Yoshioka, Keiko Takasu, Takaya Izumi, Sayaka Fujita, Daiki Nakamori, Shiori Kawasaki, Tatsuya Takahashi, Mai Yoshikawa, Koichi Ogawa, Erika Kasai

**Affiliations:** Laboratory for Drug Discovery and Disease Research, Shionogi & Co., Ltd.; Osaka, Japan; Department of Applied Pharmacology, Faculty of Pharmaceutical Sciences, University of Toyama; Toyama, Japan; R&D Supervisory Unit, Shionogi & Co., Ltd.; Osaka, Japan; Laboratory for Medicinal Chemistry Research, Shionogi & Co., Ltd.; Osaka, Japan; New Business Promotion Dept., Shionogi & Co., Ltd.; Osaka, Japan; Supply Supervisory Unit, Shionogi & Co., Ltd.; Osaka, Japan

## Abstract

Neuropathic pain remains difficult to treat effectively because of the limitations of current pain medications. The Nav1.7 sodium channel plays a crucial role in pain sensation. The development of selective inhibitors has been challenging because of the high similarity among Nav channel subtypes. To address this issue, we developed monoclonal antibodies that selectively target Nav1.7 and humanized them for clinical use for the first time. When administered systemically to neuropathic pain model rats, a potent analgesic effect was observed that lasted for at least 96 hours. Electrophysiological studies revealed that the antibody reduced mechanically-evoked and spontaneous neuronal activity in the model. Importantly, the antibodies did not impact physiological pain or motor function. Collectively, our findings suggest that these novel Nav1.7-targeting antibodies are potentially effective analgesic drugs for chronic pain, including neuropathic pain. Our novel humanized anti-Nav1.7 antibodies have the potential to be used for the development of new analgesic, and one such antibody, S-151128 is currently in clinical trial.

## Introduction

As defined by the International Association for the Study of Pain, pain refers to an unpleasant sensory and emotional experience that is associated with—or resembles that associated with— actual or potential tissue damage (Raja et al., 2020). Within this field, chronic pain is a global issue that affects a substantial portion of the population, with estimates suggesting that 10% to 55% of adults experience chronic pain (S. P. Cohen et al., 2021; De Leon-Casasola, 2013; Vellucci, n.d.).

Neuropathic pain is an important subset of chronic pain that is caused by a lesion or dysfunction in the peripheral or central nervous system (Colloca et al., 2017). Neuropathic pain is associated with greater functional impairment and lower quality of life than other types of chronic pain. This pain often leads to marked emotional distress, anxiety, and depression (Grubb, 2010; Henwood & Ellis, 2004). Furthermore, neuropathic pain is frequently characterized by ongoing or intermittent spontaneous pain, allodynia, and hyperalgesia (Finnerup et al., 2021). Treatments available for neuropathic pain include antidepressants (tricyclic antidepressants and selective noradrenaline reuptake inhibitors), gabapentinoids (pregabalin and gabapentin), topical agents (lidocaine and capsaicin), and opioids (van Velzen et al., 2020). However, although there are several pharmacological options available for managing neuropathic pain, challenges related to their efficacy, side effects, and addiction risks continue to complicate treatment strategies (Afonso et al., 2021; Fornasari, 2017; Murnion, 2018), resulting in higher unmet needs for patients experiencing neuropathic pain. There is therefore a critical need for more targeted and effective treatments that are based on the current understanding of the underlying mechanisms of neuropathic pain.

Nav1.7 is a voltage-gated sodium channel subtype that plays a crucial role in nerve conduction and transmission. It is predominantly expressed in nociceptive sensory neurons, which are responsible for detecting painful stimuli (Dib-Hajj et al., 2013; McDermott et al., 2019). Mutations in Nav1.7 (encoded by *SCN9A*) have been linked to various inherited pain syndromes, highlighting the relevance of this channel to pain sensation in humans (15,16). Gain-of-function mutations in *SCN9A* can lead to primary erythromelalgia and paroxysmal extreme pain disorder; these mutations result in lower thresholds for action potential firing, thereby causing heightened pain sensitivity and abnormal pain responses (Hameed, 2019; Huang et al., 2018). Conversely, loss-of-function mutations can lead to congenital insensitivity to pain, where individuals are unable to perceive painful stimuli because of dysfunctional Nav1.7 channels (Bennett & Woods, 2014; Cox et al., 2006). On the basis of these observations, the Nav1.7 channel is considered an attractive target for the treatment of chronic pain, including neuropathic pain.

The development of selective small-molecule inhibitors has been challenging because of the high structural similarity among the Nav subtypes (Payandeh & Hackos, 2018; Sun et al., 2014). It is difficult to achieve high selectivity for Nav1.7 over other subtypes of voltage-gated sodium channels, leading to off-target effects that can cause unwanted side effects (Hinckley et al., 2021; Naik et al., 2021; Rothenberg et al., 2019). For example, Nav1.5 is expressed in cardiac muscle and loss-of-function mutations in *SCN5A* are associated with several cardiac disorders, often resulting in life-threatening arrhythmias (Zimmer & Surber, 2008). Furthermore, Nav1.1 and Nav1.2 are predominantly expressed in the central nervous system, and mutations of these genes are associated with neurological disorders such as seizures (Eijkelkamp et al., 2012). High selectivity to the Nav1.7 subtype is therefore required for the development of safe drugs. Additionally, small molecules often face issues with metabolic stability and pharmacokinetics. Thus, ensuring that the inhibitors have a suitable pharmacokinetic profile for effective dosing without rapid degradation or clearance is essential for drug development (Safina et al., 2021). Some highly selective inhibitors, including peptide toxins targeting Nav1.7, have been reported (Murray et al., 2019; Nguyen & Yarov-Yarovoy, 2022). Nonetheless, many candidate compounds exhibit poor pharmacokinetic properties, such as high plasma protein binding and rapid clearance, which hinder their effectiveness *in vivo* (Neff & Wickenden, 2021; Nguyen et al., n.d.; Pajouhesh et al., 2020).

Numerous small-molecule drugs targeting Nav1.7 for chronic pain have failed because of side effects caused by a lack of targeting specificity or limited pharmacokinetic profiles when administered systemically (Kingwell, 2019). We therefore took an alternative approach by producing a neutralizing antibody against the Nav1.7 channel. As drugs, monoclonal antibodies have numerous benefits including strong binding affinity, high selectivity, low toxicity, and an extended half-life (Eijkelkamp et al., 2012). Despite these advantages, there is as yet no ion channel-targeting antibody that has progressed to clinical use (Eagles et al., 2022). In the present study, we generated humanized anti-Nav1.7 antibodies and demonstrated their high analgesic efficacy and long-term effectiveness in a neuropathic pain model via systemic administration. We then conducted a comprehensive evaluation of their efficacy through behavioral, electrophysiological, and histological assessments, and confirmed a lack of effects on physiological pain and motor function, with the goal of advancing toward a clinical trial for the development of a new analgesic drug.

## Results

### High binding affinity to Nav1.7 and subtype-selectivity evaluated by competitive enzyme-linked immunosorbent assay (ELISA)

The affinity and specificity of the produced novel antibodies (Clone1 and S-151128, affinity maturated variant of clone1) were confirmed using competitive ELISA. The antibodies displayed strong affinity for a Nav1.7 epitope peptide, with half-maximal inhibitory concentration (IC_50_) values of 1.52 nM (Clone1) and 0.73 nM (S-151128) in human Nav1.7 (**Fig. 1A, B, and Table 1**). In rat Nav1.7, the IC_50_ values were 2.98 nM (Clone1) and 1.51 nM (S-151128) (**Fig. 1C, D, and Table 1**). Competitive ELISA using peptides corresponding to other Nav channel subtypes revealed that the novel antibodies did not bind to any other Nav subtype (**Fig. 1A–D and Table 1**). Although the IC_50_ values for the other Nav subtypes were unable to be determined (>1000 nM), Clone1 and S-151128 exhibited selectivity for Nav1.7 of at least 650- and 1300-fold, respectively.

**Fig. 1.**
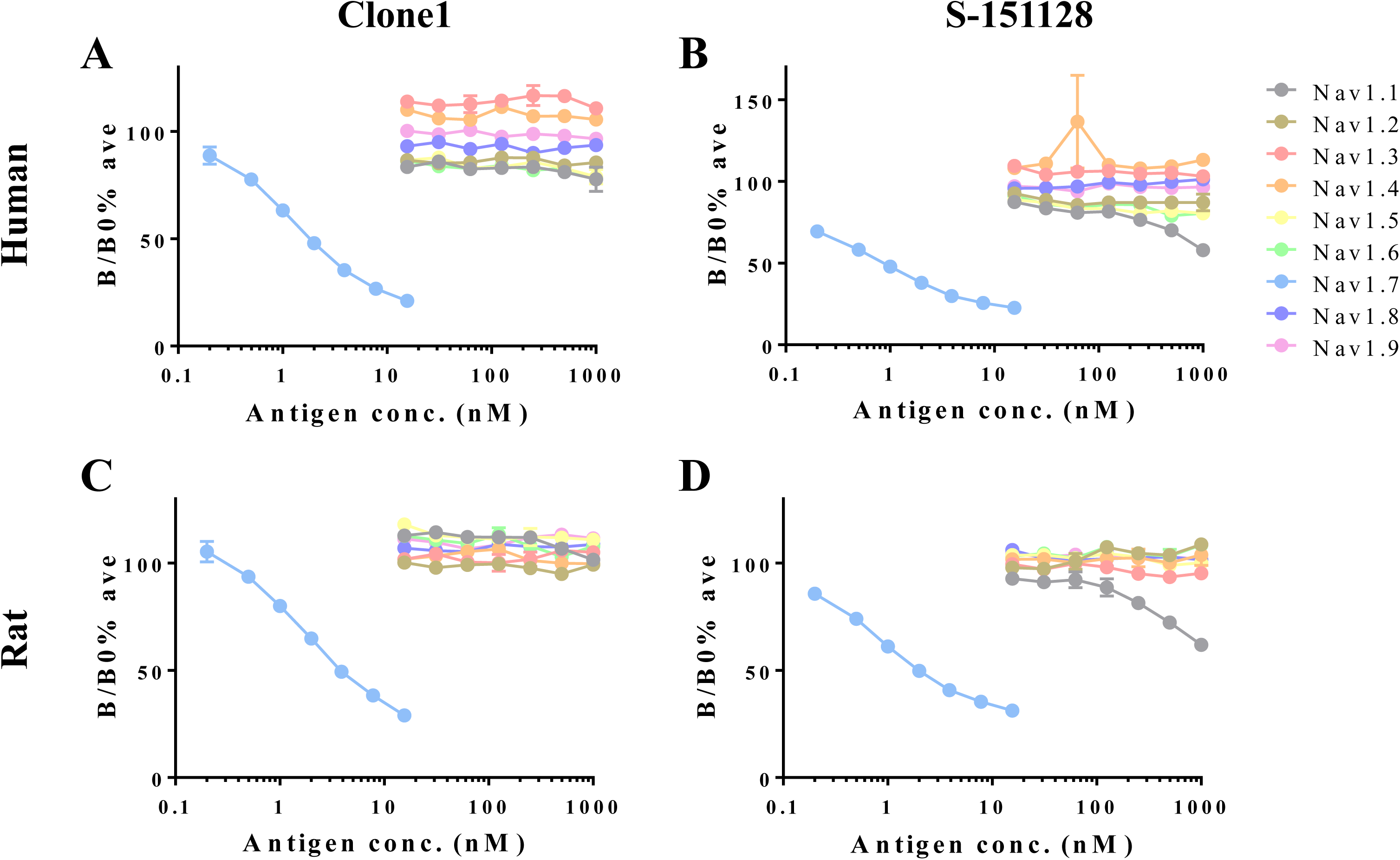
Binding affinities of clone1 and S-151128 for each Nav subtype in humans and rats. Competitive ELISA was used to evaluate the binding affinities of clone1 and S-151128 for the peptides of each Nav subtype. The inhibition ability of the antigens at varying concentrations against a constant antibody concentration of 0.7 ng/well is depicted. The binding affinities of clone1 for human (A) and rat (C) Nav subtypes are displayed. Similarly, the binding affinities of S-151128 for human (B) and rat (D) Nav subtypes are shown. Data represent the mean ± SEM from three wells; mean IC_50_ values were determined from three independent experiments.

**Table 1.** Summary of IC50 values of clone1 and S-151128 in competitive ELISA.

| Subtype<br>(Human) | Affinity (IC50, nM) |  | Subtype<br>(Rat) | Affinity (IC50, nM) |  |
| --- | --- | --- | --- | --- | --- |
|  | Clone1 | S-151128 |  | Clone1 | S-151128 |
| hNav1.7 | 1.52 | 0.73 | rNav1.7 | 2.98 | 1.51 |
| hNav1.1 | >1000 | >1000 | rNav1.1 | >1000 | >1000 |
| hNav1.2 | >1000 | >1000 | rNav1.2 | >1000 | >1000 |
| hNav1.3 | >1000 | >1000 | rNav1.3 | >1000 | >1000 |
| hNav1.4 | >1000 | >1000 | rNav1.4 | >1000 | >1000 |
| hNav1.5 | >1000 | >1000 | rNav1.5 | >1000 | >1000 |
| hNav1.6 | >1000 | >1000 | rNav1.6 | >1000 | >1000 |
| hNav1.8 | >1000 | >1000 | rNav1.8 | >1000 | >1000 |
| hNav1.9 | >1000 | >1000 | rNav1.9 | >1000 | >1000 |

### Binding of antibodies to Nav1.7 expressed in human embryonic kidney 293 (HEK) cells

To confirm the binding of the antibodies to Nav1.7 on the cell membrane, immunocytochemistry was performed using HEK cells stably expressing human or rat Nav1.7. Our antibodies were used as the primary antibody, and their binding to Nav1.7 was visualized using a fluorescence-conjugated secondary antibody. As depicted in **Fig. 2**, fluorescence derived from Nav1.7 was observed in the presence of each primary antibody, whereas the negative control group (lacking the primary antibody) displayed minimal fluorescence. These results indicate that our novel antibodies can bind to Nav1.7 on the cell membrane.

**Fig. 2.**
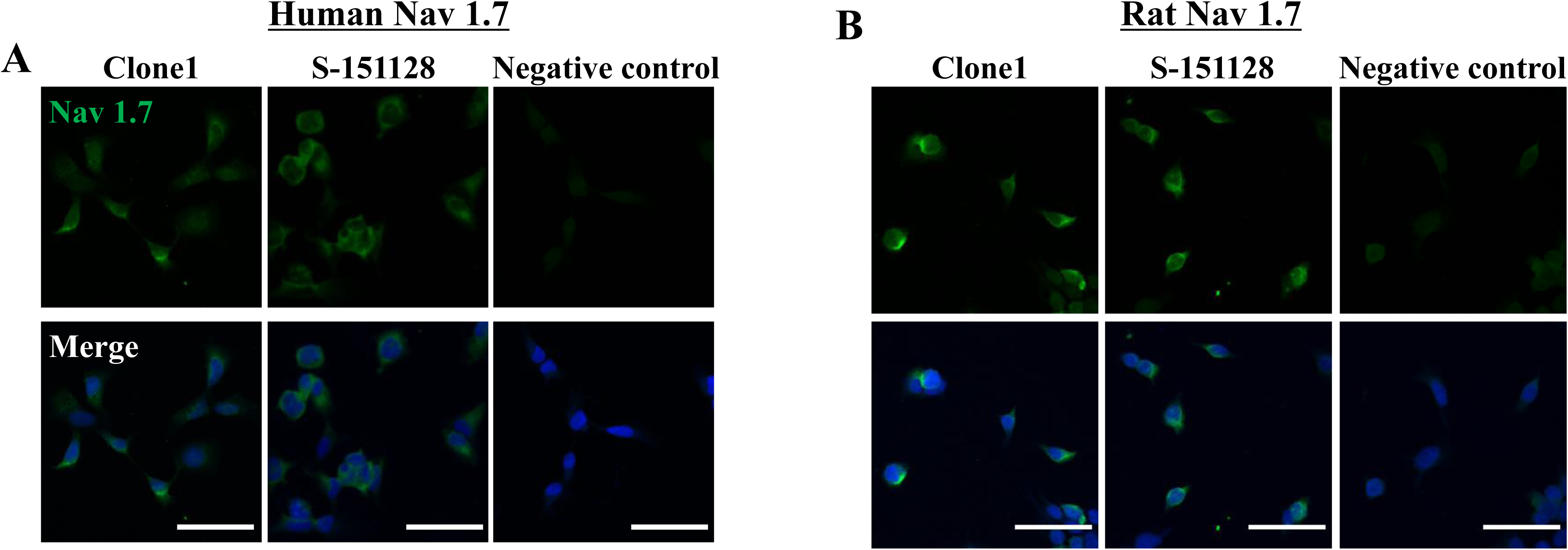
Immunofluorescent staining of HEK cells expressing Nav subtypes using the antibodies. Representative images of HEK cells stained with the novel antibodies as the primary antibody (green) and a nuclear stain (blue) are shown. HEK cells stably expressing human Nav1.7 are depicted in (A), and those expressing rat Nav1.7 are shown in (B). Negative control images, where the primary antibody was not included, displayed little specific staining. Scale bar = 50 µm.

### Functional inhibition of Nav1.7 expressed in HEK cells and rat dorsal root ganglion (DRG) neurons

To evaluate the functional inhibition of our antibodies on Nav1.7 channels, patch-clamp recordings were conducted with a whole-cell voltage-clamp configuration using HEK cells stably expressing human/rat Nav1.7 channels. The perfusion of each antibody (100 µg/mL) for 10 minutes resulted in significantly lower peak sodium currents compared with the negative control antibody (**Fig. 3A–F and Fig. S1**). The percentage inhibitions of sodium currents by clone1 and S-151128 were 21.4/22.2% (human/rat Nav1.7) and 24.2/16.7% (human/rat Nav1.7), respectively. Although the inhibitory effect on sodium currents was partial, previous reports have demonstrated that even the partial inhibition of sodium currents significantly inhibits neuronal action potentials (Yau et al., 2010). Given that neuronal activity plays a critical role in transmitting pain signals from the periphery to the central nervous system (Goodwin & McMahon, 2021; Moldovan et al., 2013), we investigated the effects of our antibodies on neuronal action potentials in rat DRG neurons. Exposure to each antibody at 100 µg/mL significantly reduced neuronal action potentials (**Fig. 3G–J**), suggesting that these antibodies may attenuate pain signals *in vivo*.

**Fig. 3.**
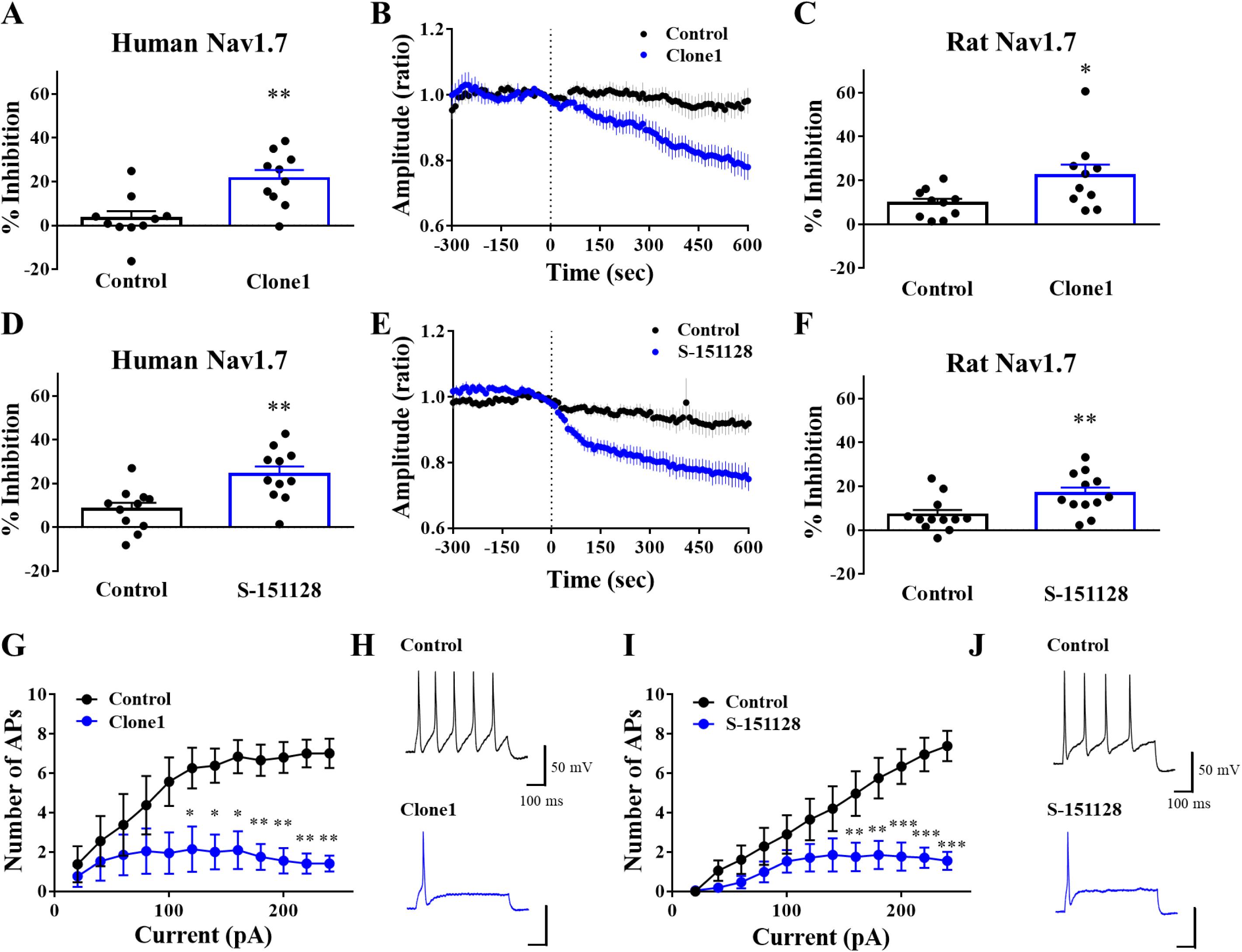
Functional inhibition of the antibodies with *in vitro* electrophysiology. Whole-cell patch-clamp recordings were conducted on HEK cells expressing human Nav1.7 to assess the inhibitory effects of the antibodies at concentrations of 100 µg/mL on the sodium current (A–F). Data are presented as the mean ± SEM. Significance was determined using a *t*-test; \**p* < 0.05, \*\**p* < 0.01 compared with the control group; *n* = 10–12. The time-courses of peak currents during the experiments are shown in (B) and (E); the dotted lines indicate the start of antibody perfusion. To evaluate the number of APs, membrane potential recording was performed on visually identified rat DRG neurons (G and H). Data are presented as the mean AP ± SEM for each current injection. Representative traces of APs induced by 140 pA current injection are shown in H and J. Significance was determined using two-way ANOVA followed by the Holm–Šidák test; \**p* < 0.05, \*\**p* < 0.01, \*\*\**p* < 0.01 compared with the control group; *n* = 7–8. The negative control antibody was used as the control in all experiments. AP: action potential.

### Antibodies increase the paw withdrawal threshold (PWT) in a partial sciatic nerve ligation (PSNL) model

We next investigated the efficacy of intravenous Nav1.7 antibody administration as a potential therapeutic drug for pain relief. Previous studies have reported the analgesic effects of anti-Nav1.7 antibodies when administered intrathecally or into the DRG (Bang et al., 2018; Xia et al., 2016). However, in terms of practicality and ease of clinical use, intravenous antibody administration would be more convenient. We therefore aimed to evaluate the efficacy of intravenous antibody administration. To assess the effects of our antibodies on pain behavior *in vivo*, we used a rat model of PSNL and von Frey filaments (vFF). Systemic antibody administration via an intravenous route at various doses (Clone1: 0.5, 1.5, 5, or 15 mg/kg; S-151128: 0.03, 0.1, 0.3, 1, 3, or 10 mg/kg) resulted in dose-dependent increases in the PWT, indicating reduced pain sensitivity (**Fig. 4A, B**). The 0.5 mg/kg dose of clone1 exhibited efficacy similar to that of pregabalin (10 mg/kg), with the peak efficacy observed at 5 mg/kg. Similarly, the 0.1 mg/kg dose of S-151128 displayed an efficacy similar to that of pregabalin, reaching its maximum efficacy at 3 mg/kg. Importantly, even 96 hours after administration, the efficacies of both antibodies remained superior to that of pregabalin. Plasma/serum concentrations of the antibodies were measured at all timepoints and the doses were evaluated using behavioral testing. The effects of the antibodies on the PWT were highly correlated with their plasma/serum concentrations (adjusted *r*^2^ = 0.880 in clone1 and 0.939 in S-151128) (**Fig. 4C, D**).

**Fig. 4.**
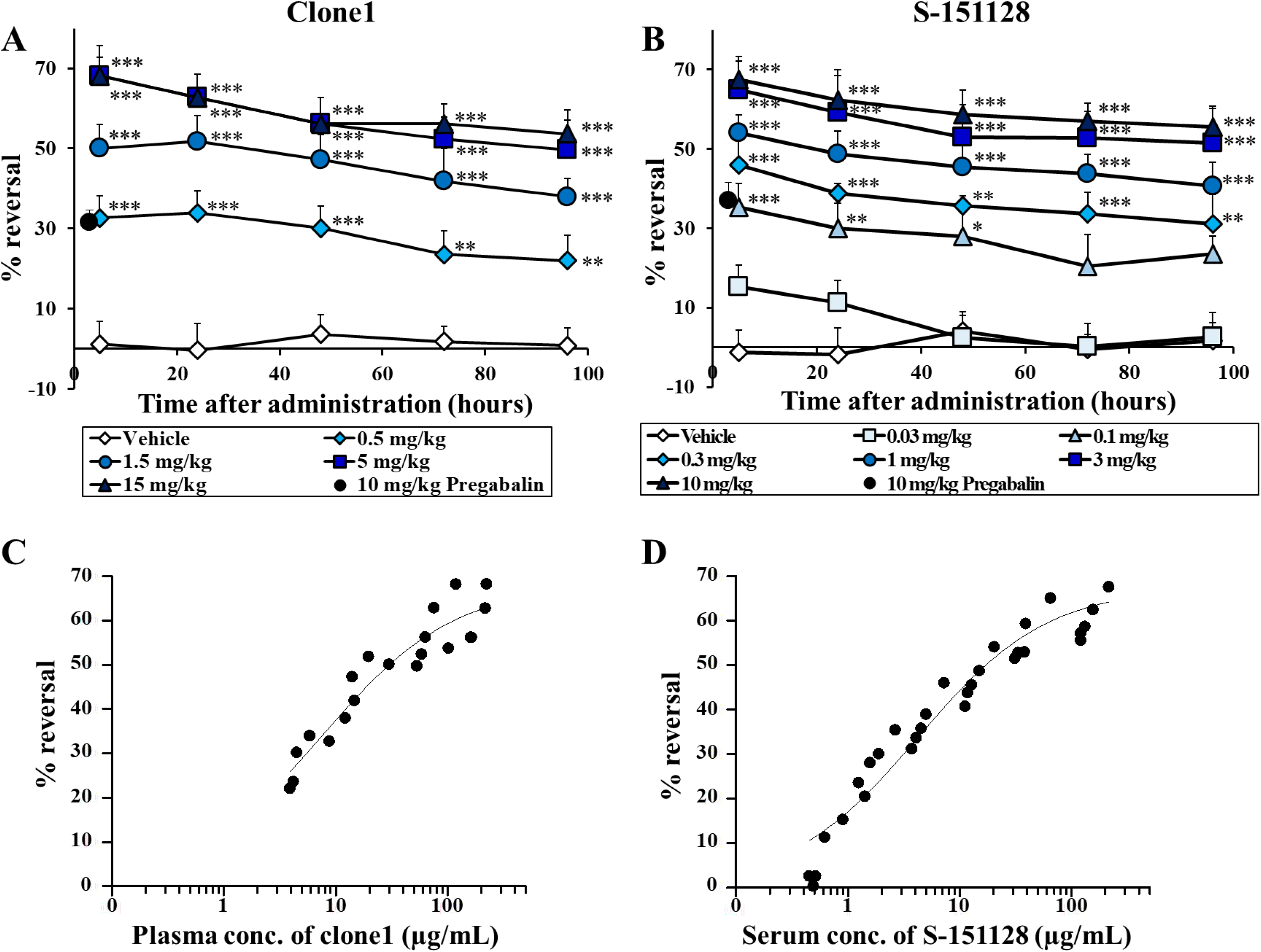
Effects of the antibodies on the PWT in PSNL model rats. The analgesic effects of clone1 (A) and S-151128 (B) were evaluated over time. The PWT was measured before treatment and at 5, 24, 48, 72, and 96 hours after the single administration of each antibody. Data are presented as the mean ± SEM, with a sample size of 8–10. Significance was determined using a two-way ANOVA followed by Dunnett’s *post hoc* test; \**p* < 0.05, \*\**p* < 0.01, and \*\*\**p* < 0.001 compared with the vehicle-treated group at the respective timepoints after treatment. The relationship between the efficacy and plasma/serum concentration of each antibody is shown (C and D). Each closed circle represents the efficacy and plasma/serum concentration at all doses and timepoints for testing pain behavior.

### Suppression of neural activity in *in vivo* extracellular recordings

To further assess the pharmacological effects of our antibodies, we conducted electrophysiological studies. Extracellular recordings of action potentials from spinal dorsal horn neurons were performed 2 weeks after PSNL induction to examine both mechanically evoked and spontaneous excitatory input from peripheral afferents to the spinal cord (**Fig. 5A**). The electrode was implanted at a depth of 20–150 µm from the surface; this depth did not differ between the sham and PSNL groups (**Fig. 5B**). The spontaneous firing rate (without stimulation) was significantly higher in the PSNL model (**Fig. 5C, D**). Similarly, the PSNL model exhibited significantly higher neural activity in response to mechanical stimulation by vFF compared with the sham group (**Fig. 5E, F**). These results are consistent with previous reports of neuropathic pain models (Gong et al., 2019; Uta et al., 2019). We then evaluated the inhibitory effects of clone1 on neural activity in the PSNL model. Both vFF-induced and spontaneous neural activity were evaluated 5–8 hours after the intravenous injection of clone1 at doses at which the antibody showed maximal efficacy in the behavioral test (0.5, 5, or 15 mg/kg,). The electrode depth did not differ between the vehicle and clone1-treated groups (**Fig. 6A**). Clone1 led to significant dose-dependent decreases in both spontaneous and vFF-evoked firing (**Fig. 6B–E**). The increased PWT and the inhibition rate of neural firing at each dose of clone1 are summarized in **Fig. 6F**. Both the behavioral and electrophysiological evaluations demonstrated consistent efficacy, thus supporting the effectiveness of this antibody.

**Fig. 5.**
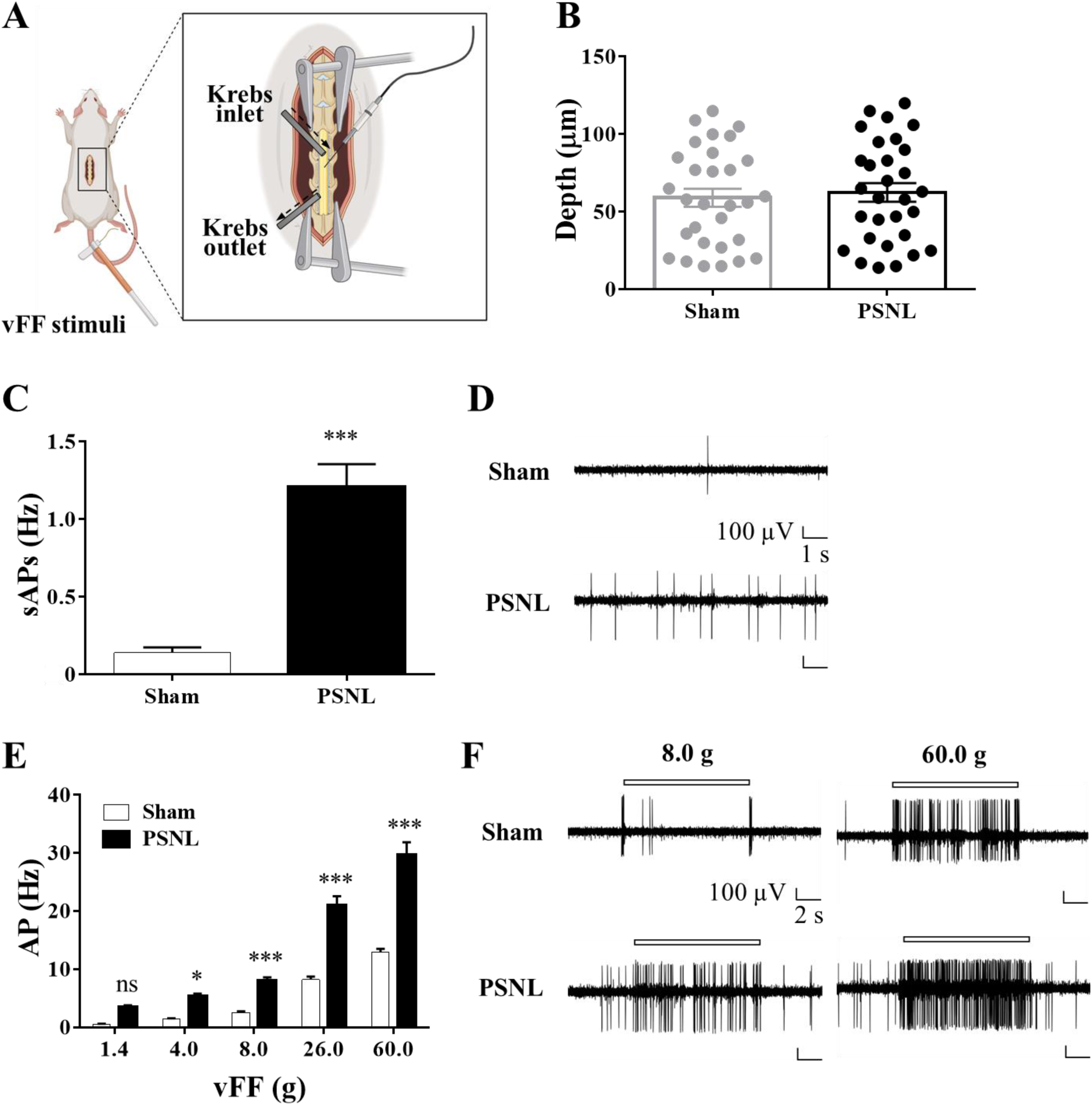
Increased activity of superficial dorsal horn neurons in PSNL model rats. The experimental set-up is presented in (A). Rats were fixed with stereotaxic instruments under urethane anesthesia, and an electrode was inserted into superficial dorsal horn. Mechanical stimuli were administered to the identified RF with vFF. The depth of electrode placement from the surface of the spinal cord is shown in (B). Data were collected from five neurons each from six rats. A *t-*test indicated no significant difference between the groups. The spontaneous discharge rates in both the sham and PSNL rats are illustrated in (C). Significance was determined using a *t*-test; \*\*\**p* < 0.001 compared with the sham group (*n* = 30 from six rats). The analysis of vFF-evoked AP frequency is shown in (E). vFF at 1.4, 4, 8, 26, and 60 g were used for the analysis, with a sample size of 30 (from six rats). Significance was determined using the Holm–Šidák test; \**p* < 0.05, \*\**p* < 0.01, and \*\*\**p* < 0.001 compared with the sham group. Representative traces of APs recorded from sham and PSNL rats with/without vFF are shown in (D and F). All data are presented as the mean ± SEM. AP: action potential, ns: not significant, RF: receptive field, sAP: spontaneous action potential.

**Fig. 6.**
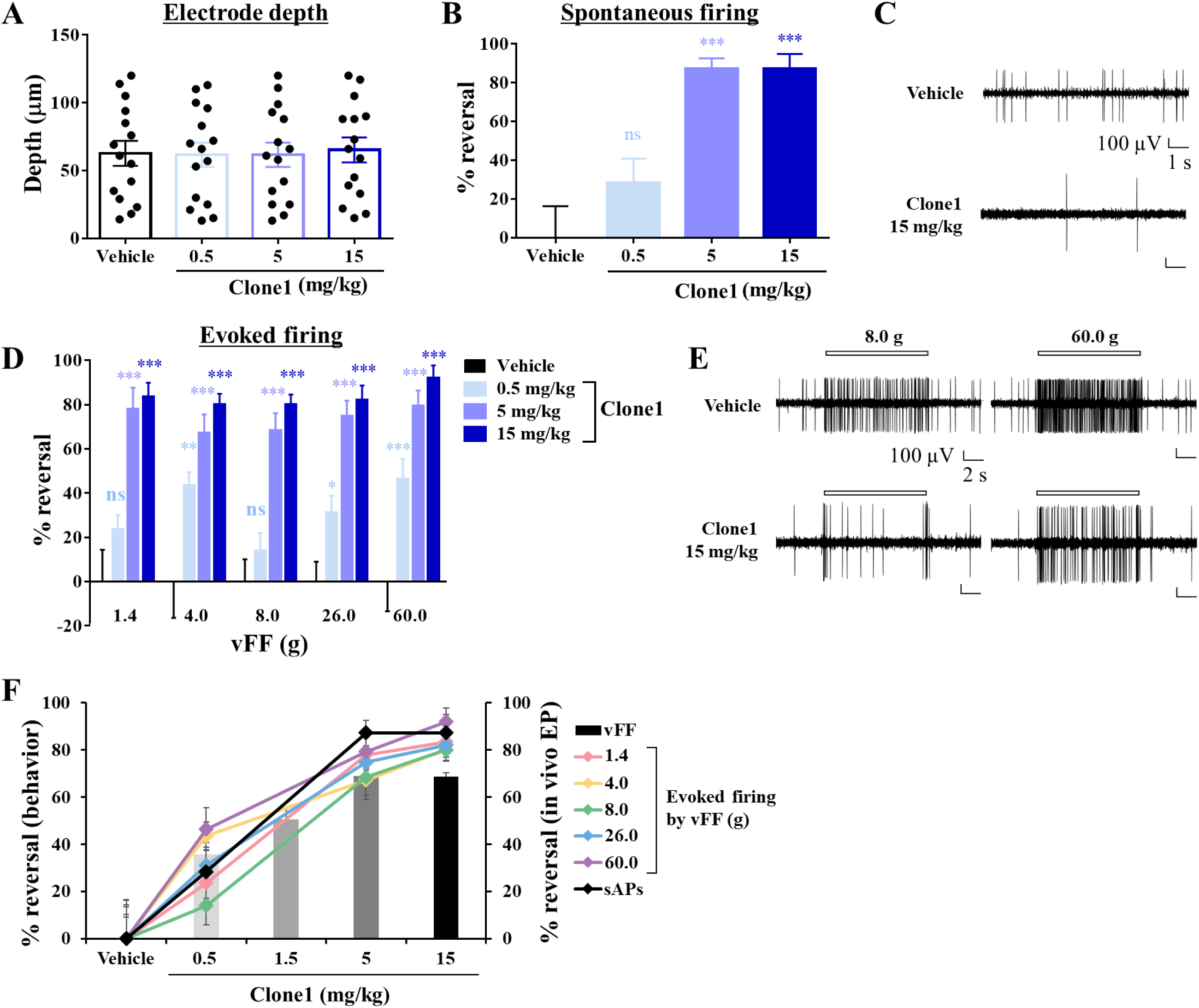
Effects of clone1 on the activity of dorsal horn neurons. The depth of electrode placement from the surface of the spinal cord is shown in (A). Data were collected from five neurons each from three rats. Data are presented as the mean ± SEM. Dunnett’s test indicated no significant difference between the groups. Two weeks after the PSNL operation, the PSNL rats received intravenous injections of clone1 (0.5, 5, or 15 mg/kg) or vehicle. Neuronal activity was recorded 5–8 hours after clone1 injection. Data showing spontaneous activity (B and C) and evoked firing (D and E) are presented as the mean ± SEM, *n* = 15 (from three rats). Statistical analysis was performed using the Holm–Šidák test; \**p* < 0.05, \*\**p* < 0.01, and \*\*\**p* < 0.001 compared with the vehicle group. Representative traces of APs are shown in (C and E). All pharmacological data from the behavioral testing and electrophysiological experiments are summarized in (F). EP: electrophysiological test, ns: not significant, sAP: spontaneous action potential.

### Inhibition of phosphorylation of extracellular signal-regulated kinase (ERK) in rat DRG neurons

The phosphorylation of ERK in DRG neurons is reportedly augmented in neuropathic pain models (Ma & Quirion, 2005; Obata et al., 2004). Furthermore, ERK in DRG neurons undergoes phosphorylation in response to painful stimuli, and this phosphorylation is considered to reflect pain signals (Gao & Ji, 2009). In the present study, we therefore aimed to confirm the inhibitory effect of clone1 on pain signals by immunohistochemically assessing phosphorylated (p)ERK in the DRG after vFF stimulation in the PSNL model. vFF (60 g) prompted ERK phosphorylation in DRG neurons, with notably higher levels observed in the PSNL model than in sham rats. Importantly, clone1 administration significantly reduced the number of pERK-positive cells (**Fig. 7A, B**).

**Fig. 7.**
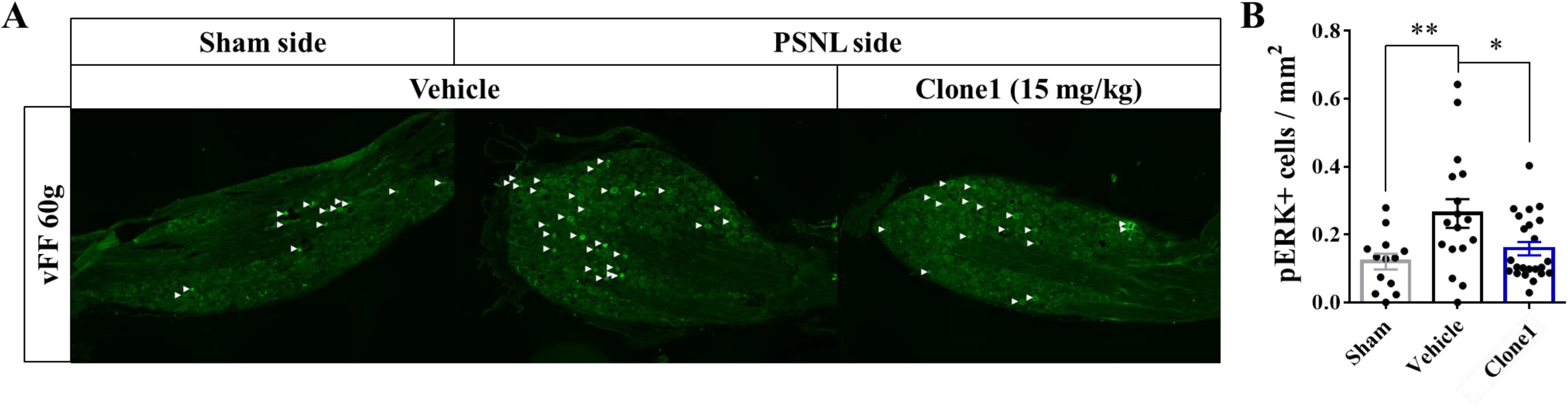
Effects of clone1 on vFF-evoked ERK phosphorylation in DRG neurons. Representative images of the rat DRG stained with anti-pERK antibody (green) are shown in (A). Arrowhead indicates pERK-positive cells. The numbers of pERK-positive cells were counted in the sham, vehicle-treated PSNL, and clone1-treated PSNL model rats (B). Data are presented as the mean ± SEM (sham: *n* = 15, vehicle: *n* = 16, clone1: *n* = 23, *n* = slice, 2–4 slices per rat). Statistical analysis was performed using one-way ANOVA with Dunnett’s test; \**p* < 0.05 and \*\**p* < 0.01 compared with the vehicle group.

### Effects of the antibody on physiological pain and motor function

Given that loss-of-function mutations in Nav1.7 result in congenital insensitivity to pain (Bennett & Woods, 2014; Cox et al., 2006), we explored the effects of the antibody on pain behaviors and signals under physiological conditions. To evaluate the effects of the antibody on physiological pain behavior, the PWT of sham-side hindlimbs were assessed 5 hours after the administration of each antibody. In contrast to their effects in hindlimbs on the nerve-ligated side, the antibodies had no effects on the PWT in hindlimbs on the sham side (**Fig. 8A, B**). We next assessed neuronal activity in sham rats following clone1 administration. Notably, clone1 had no effects on spontaneous neuronal action potentials or vFF-evoked neuronal activity in sham rats (**Fig. 8C, D**).

**Fig. 8.**
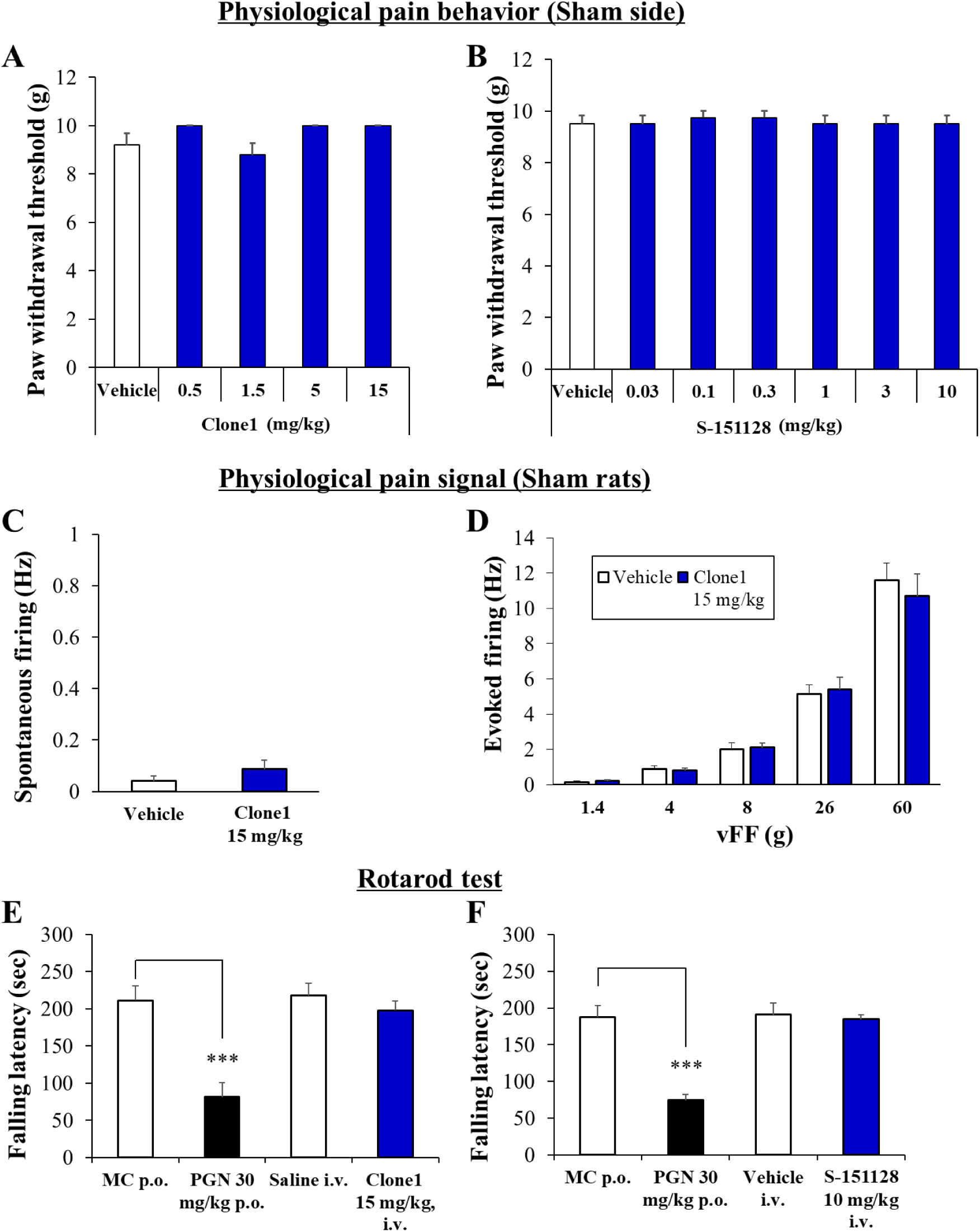
Effects of the antibodies on physiological pain signals and motor function. Rats received intravenous injections of clone1, S-151128, or vehicle for the assessment of physiological functions. The PWT of the sham side was evaluated 5 hours after the administration of clone1 (0.5, 1.5, 5, or 15 mg/kg), S-151128 (0.03, 0.1, 0.3, 1, 3, or 10 mg/kg), or vehicle. *n* = 5 for clone1 and *n* = 8 for S-151128. Dunnett’s test indicated no significant difference between the treatment and vehicle groups. Neuronal activity was recorded 5–8 hours after clone1 injection. *n* = 15 (from three rats). The *t*-test and Holm–Šidák test indicated that there were no significant differences between the groups in terms of spontaneous firing and evoked firing, respectively. Normal rats received intravenous injections of clone1 (15 mg/kg) or vehicle for the rotarod test. Falling latency was measured 5 hours after clone1 injection. PGN was used as a positive control, and MC was used as the control for PGN. *n* = 8–9. No significant difference was observed between clone1 and vehicle (intravenous). All data are presented as the mean ± SEM. Statistical analysis using the *t*-test revealed a significant reduction by PGN compared with oral MC; \*\*\**p* < 0.001. PGN: pregabalin, MC: 0.5% methyl cellulose.

Commonly used analgesics, such as opioids and pregabalin, have side effects such as sedation and dizziness, which can impair motor function. These adverse effects often restrict their use in specific clinical situations (Rissardo & Caprara, 2020; Spahn et al., 2018). To evaluate the effects of anti-Nav1.7 antibody on motor function, we conducted the rotarod test after intravenous antibody administration. As previously reported (Khan et al., 2018; Yokoyama et al., 2007), the positive control (pregabalin, 30 mg/kg) significantly reduced the time spent on the rotarod, indicating impaired motor function. By contrast, antibody administration had no effect on the time spent on the rod, suggesting that our antibodies do not impair motor function (**Fig. 8E, F**).

## Discussion

We have developed novel humanized monoclonal antibodies against the Nav1.7 sodium channel. These antibodies were able to specifically bind Nav1.7 and inhibit its function *in vitro*. Notably, the antibodies demonstrated significant and long-lasting analgesic effects in a rat model of neuropathic pain but had no effects on physiological pain and caused no motor function impairment. The analgesic effects were evaluated by not only behavioral assessments but also electrophysiological and immunohistochemical evaluations.

In the current study, we demonstrated the efficacy of our antibodies via systemic intravenous administration in a rat neuropathic pain model. Although previous reports have demonstrated the analgesic effects of antibodies against Nav1.7, they have only evaluated the efficacy of intrathecal administration or local administration into the DRG rather than systemic administration (Bang et al., 2018; Martina et al., 2024; Xia et al., 2016). In clinical practice, the local administration of anesthetics (e.g. lidocaine) is performed for nerve block or spinal block therapy; however, this method has various disadvantages. For example, it has as a short duration of efficacy, is time-consuming, causes temporary motor weakness and numbness, and there is a risk of infection at the injection site (Kent & Bollag, 2010; Varitimidis et al., 2009; Wiegel et al., 2007). Hence, there is a pressing need for more convenient analgesic therapies. In the present study, we provided evidence for the potent efficacy of Nav1.7 antibodies even when administered systemically. Notably, this analgesic effect persisted for up to 96 hours, surpassing the efficacy of pregabalin (used as a control drug). To the best of our knowledge, no previous reports have documented such sustained and pronounced efficacy through the systemic administration of Nav1.7 antibodies. The reported half-life of humanized antibodies in rodents ranges from several to approximately 10 days (Espinoza et al., 2019; Martin et al., 2010; Webster et al., 2023), suggesting that prolonged exposure may have contributed to the observed long-term efficacy. Furthermore, given that the half-life of humanized antibodies is generally longer in humans than in rodents (primarily because of disparities in the binding affinity of the neonatal Fc receptor between species (Proetzel & Roopenian, 2014)), it is reasonable to expect even greater long-term efficacy in humans.

Our novel antibodies inhibited vFF-induced paw withdrawal and neuronal firing as well as spontaneous neuronal firing. Increased responses to vFF-induced innocuous stimuli in terms of both behavior and neuronal firing are believed to reflect mechanical allodynia (Barrot, 2012; Pitcher & Henry, 2004). It has also been reported that neuronal firing in C fibers is associated with spontaneous pain (Djouhri et al., 2006; Kleggetveit et al., 2012). In the current study, we evaluated neural activity in a superficial region of the spinal cord (20–150 µm from the spinal cord surface, lamina II), which is an area of C fiber input (D’Mello & Dickenson, 2008), suggesting that our findings reflect C fiber activity. As shown in **Fig. 6**, the inhibitory effects were similar for all doses in each evaluation, indicating that our antibody can suppress both mechanical allodynia and spontaneous pain at the same dose. Allodynia and spontaneous pain are considered to be the main symptoms of neuropathic pain (Finnerup et al., 2021). These findings therefore suggest that our antibodies may effectively inhibit both spontaneous pain and allodynia in patients with neuropathic pain.

Our antibodies demonstrated partial inhibition in the *in vitro* evaluation of sodium currents (shown in **Fig. 3**). The antibodies bind to the E3 loop in domain III of the Nav1.7 structure. It has been reported that many ion channel antibodies target this E3 region and can significantly, but not completely, inhibit currents (Gómez-Varela et al., 2007; Naylor & Beech, 2009; Xu et al., 2005). Although the exact mechanism underlying this effect remains unclear, various possibilities have been proposed, including partial occlusion of the ion channel pore, allosteric modulation via the E3 region, and channel internalization (Naylor & Beech, 2009). Sodium channel blockers such as flufenamic acid can partially inhibit voltage-gated sodium currents in hippocampal neurons; however, this partial inhibition reduces neuronal firing, indicating that the partial inhibition of sodium currents can effectively suppress neural excitability (Yau et al., 2010). In the present study, even with the antibody-induced partial inhibition of sodium currents, neuronal firing in the rat DRG was significantly suppressed (**Fig. 3**). This observed inhibitory effect was similar to that observed in neurons derived from induced pluripotent stem cells obtained from patients with congenital insensitivity to pain, as well as in neurons with Nav1.7 knockout (McDermott et al., 2019). Additionally, our antibody demonstrated adequate efficacy in *in vivo* behavioral and electrophysiological evaluations (**Figs. 4, 6**). Together, these results suggest that even with the partial inhibition of sodium currents, our antibody exhibits marked functional effects.

Antibodies typically have limited penetration into peripheral nerve tissue, primarily because of biological barriers such as the blood–nerve barrier (BNB) (Liu et al., 2018; Weerasuriya & Mizisin, 2011). In sham rats, our antibody did not exhibit any effects on neuronal firing and motor function (**Fig. 8**). However, BNB disruption in neuropathic pain models may lead to the enhanced tissue penetration—and therefore therapeutic effects—of antibodies (Richner et al., 2019). In rodent models of neuropathic pain, such as PSNL and chronic constriction injury, there is clear evidence that the BNB becomes impaired (Lim et al., 2014; Moreau et al., 2016; Reinhold et al., 2018). Similarly, in a diabetic neuropathy model, both the BNB and blood–brain barrier are reportedly disrupted (Ben-Kraiem et al., 2091; Huber et al., 2006; Salameh et al., 2016). The BNB is also impaired in patients with diabetic neuropathy, allowing for the increased penetration of immunoglobulin G (IgG) into nerve tissue (Poduslo et al., 1988). Collectively, these findings suggest that systemically administered antibodies may be able to reach nerve tissue and exert therapeutic effects in patients with neuropathic pain, such as diabetic neuropathy.

Our study has some limitations. For example, the tissue concentrations of the antibodies were not measured. Although we attempted to quantify the antibody concentration in the sciatic nerve, the levels were unable to be accurately assessed. Consequently, it remains unknown whether the antibody directly interacts with the sciatic nerve and contributes to the observed effectiveness. Furthermore, efficacy data were only presented for the PSNL model in the present study. Although previous reports suggest that Nav1.7 inhibition demonstrates efficacy against neuropathic pain, further investigations using our antibodies are warranted to validate this assumption. In addition, our electrophysiological experiments evaluated the effects of the antibodies to approximately 5 hours post-administration, with peak analgesic effects at this timepoint. However, the extension of neural activity suppression beyond this timeframe remains uncertain. Finally, the absence of a comparison between pre- and post-treatment conditions introduces uncertainty regarding the inhibition of recorded neurons by the antibody. Nevertheless, a clear distinction between the vehicle and treatment group was evident, suggesting efficacy.

Monoclonal antibodies are a rapidly growing field for analgesia. Tumor necrosis factor alpha antibodies, also known as tumor necrosis factor inhibitors, are used as standard care for various autoimmune diseases and can provide pain relief in certain conditions, particularly those involving chronic inflammation (Monaco et al., 2015). Calcitonin gene-related peptide antibodies are also increasingly recognized for their role in pain relief, particularly in the context of migraines and cluster headaches (F. Cohen et al., 2022). Although Nav1.7 is a promising drug target, there are currently no therapeutic antibodies against this ion channel. Our work demonstrates the feasibility of therapeutic antibody development for the treatment of ion channel-related diseases. Our antibodies selectively bound to Nav1.7 and inhibited Nav1.7 channel function *in vitro*; they also had long-term *in vivo* analgesic effects in an animal neuropathic pain model without affecting physiological pain or motor function. Furthermore, these antibodies were humanized from mouse antibodies. Antibody humanization is a critical process in the development of therapeutic antibodies. This process offers several important advantages that are primarily aimed at enhancing the safety and efficacy of these treatments (Kuramochi et al., 2019; Ling et al., 2021). Thus, these humanized monoclonal antibodies may confer many therapeutic advantages including selectivity, duration, and safety. These antibodies have the potential to be used for the development of new therapeutics against pain, and one such antibody, S-151128 is currently in clinical trial.

## Materials and methods

### Study design

The aim of the present study was to analyze the pharmacological effects of the newly developed anti-Nav1.7 antibodies, with the goal of conducting clinical trials in the future. To achieve this, we first humanized the antibodies to make them suitable for human administration. We hypothesized that the antibodies would have high selectivity to Nav1.7 and exhibit strong therapeutic effects, and conducted non-clinical *in vitro*/*in vivo* efficacy evaluations. Additionally, we conducted evaluations of the long-term pharmacological effects of administration because we expected the antibodies to demonstrate prolonged efficacy. Furthermore, we confirmed the therapeutic effects of the antibodies from multiple perspectives including behavioral evaluations, the inhibition of neuronal activity using electrophysiological methods, and the histological evaluation of neuronal activity markers. Sprague Dawley rats and cultured DRG cells from rats were used. Sample sizes and experimental designs were determined on the basis of previously published data from our laboratories or similar experiments in the field. The exact numbers (*n*) used in each study are indicated in the respective figure legends. Experiments were completed over multiple time periods to ensure that replication was observed. All animals were randomly assigned to the experimental and control groups, and the experimenters were blinded for the behavioral testing.

### Generation of anti-Nav1.7 antibodies

We selected a peptide corresponding to the domain III, E3 extracellular loop C-terminus region of human Nav1.7 (UniProtKB/Swiss-Prot: Q15858) as an antigen. The peptide (1424-QPKYEYSL-1431) was synthesized by introducing a Cys residue at the N-terminus (manufactured by Toray Industries, Otsu, Japan), conjugated to keyhole limpet hemocyanin (KLH) as an immunogen. Next, 4- to 6-week-old female A/J Jms Slc mice were injected intraperitoneally with 0.1 mg of KLH-conjugated peptide (KLH-CQPKYEYSL) emulsified in complete Freund’s adjuvant (Difco/Becton Dickinson, Franklin Lakes, NJ). Four additional injections of 0.1 mg of KLH-conjugated peptide emulsified in incomplete Freund’s adjuvant followed at 3-week intervals. Eight days after the fifth immunization, mice were injected with 0.1 mg of adjuvant-free antigen. Three days after the final injection, splenocytes from the immunized mice were collected aseptically and fused to murine myeloma cells using 50% (weight/volume) polyethylene glycol 4000. Human Nav1.7 binding antibodies were then identified by performing ELISA using hybridoma culture supernatants against the immunogenic peptide.

### Competitive ELISA

First, 96-well MaxiSorp plates were coated with 10 µg/mL of anti-human IgG Fc antibody in 50 mM Tris-HCl (pH = 7.5) by overnight incubation at room temperature. Next, plates were washed with wash buffer (Nacalai Tesque, Kyoto, Japan) before being blocked with blocking buffer containing 4% Block ACE and 10% sucrose for 2 hours at room temperature. After a washing step, 50 μL of antibodies, 50 µL of biotinylated peptide (0.7 ng/well, each peptide sequence is shown in Table 2), 50 µL of competitor (non-labeled) peptide, and streptavidin–horseradish peroxidase (45 ng/well) were added and incubated overnight at 4°C. After another wash, the plates were incubated with 100 µL of the substrate solution tetramethylbenzidine. After adequate enzymatic reaction, the reaction was stopped with 100 µL of 0.5 M sulfuric acid. Absorbance values were measured at 450 nm using a SpectraMax Paradigm (Molecular Devices, San Jose, CA). The IC_50_ value indicates the antigen concentration when the inhibition rate reaches 50%. All experiments were performed in triplicate.

**Table 2.** Sequence of biotinylated peptides.

| Subtype | Biotinylated peptides |
| --- | --- |
| Human Nav1.1 | Cys(Biotinyl-PEG2-maleimide)-SRNVELQPKYEESL-NH2 |
| Human Nav1.2 | Cys(Biotinyl-PEG2-maleimide)-SRNVELQPKYEDNL-NH2 |
| Human Nav1.3 | Cys(Biotinyl-PEG2-maleimide)-SRDVKLQPVYEENL-NH2 |
| Human Nav1.4 | Cys(Biotinyl-PEG2-maleimide)-SREKEEQPQYEVNL-NH2 |
| Human Nav1.5 | Cys(Biotinyl-PEG2-maleimide)-SRGYEEQPQWEYNL-NH2 |
| Human Nav1.6 | Cys(Biotinyl-PEG2-maleimide)-SRKPDEQPKYEDNI-NH2 |
| Human Nav1.7 | Cys(Biotinyl-PEG2-maleimide)-SVNVDKQPKYEYSL-NH2 |
| Human Nav1.8 | Cys(Biotinyl-PEG2-maleimide)-SREVNMQPKWEDNV-NH2 |
| Human Nav1.9 | Cys(Biotinyl-PEG2-maleimide)-STEKEQQPEFESNS-NH2 |
| Rat Nav1.1 | Cys(Biotinyl-PEG2-maleimide)-SRNVELQPKYEESL-NH2 |
| Rat Nav1.2 | Cys(Biotinyl-PEG2-maleimide)-SRNVELQPKYEDNL-NH2 |
| Rat Nav1.3 | Cys(Biotinyl-PEG2-maleimide)-SRDVKLQPIYEENL-NH2 |
| Rat Nav1.4 | Cys(Biotinyl-PEG2-maleimide)-SREKEEQPHYEVNL-NH2 |
| Rat Nav1.5 | Cys(Biotinyl-PEG2-maleimide)-SRGYEEQPQWEDNL-NH2 |
| Rat Nav1.6 | Cys(Biotinyl-PEG2-maleimide)-SRKPDEQPDYEGNI-NH2 |
| Rat Nav1.7 | Cys(Biotinyl-PEG2-maleimide)-SVNVNEQPKYEYSL-NH2 |
| Rat Nav1.8 | Cys(Biotinyl-PEG2-maleimide)-SGEINSQPNWENNL-NH2 |
| Rat Nav1.9 | Cys(Biotinyl-PEG2-maleimide)-SREKDEQPDFEANL-NH2 |

### Antibody humanization

A humanized antibody with activity equivalent to the mouse antibody, selected through hybridoma screening, was generated by complementarity-determining region grafting and the introduction of multiple back-mutations.

### Cells

HEK cells expressing human Nav1.7 sodium channels were obtained from Charles River Laboratories (Wilmington, MA). To generate a HEK cell line expressing rat Nav1.7 sodium channels, we transfected the cells with a plasmid encoding rat Nav1.7. Subsequently, a single clone was selected and maintained for all further experiments (immunocytochemistry and electrophysiology).

### Immunocytochemistry

HEK cells expressing human/rat Nav1.7 were fixed in 4% paraformaldehyde for 10 minutes before being blocked with 3% bovine serum albumin for 60 minutes at room temperature. The cells were then incubated in our antibodies (100 µg/mL) overnight at 4°C. After washing with phosphate-buffered saline (PBS), secondary antibodies (mouse anti-human IgG, 31137, Thermo Fisher Scientific, San Jose, CA) were added at 10 µg/mL and incubated overnight at 4°C. After washing with PBS, tertiary antibodies (donkey anti-mouse IgG Alexa 488, A21202, Thermo Fisher Scientific) were added at 2 µg/mL and incubated for 60 minutes at room temperature. Coverslips were then mounted using VECTASHIELD with 4’,6-diamidino-2-phenylindole (Vector Laboratories, Burlingame, CA), and the immunostaining was visualized using a BZ-X710 microscope (KEYENCE, Osaka, Japan).

### Whole-cell voltage-clamp recording in HEK cells

Whole-cell voltage-clamp recordings were performed at room temperature using a Double Integrated Patch Amplifier (Sutter Instrument, Novato, CA) controlled with an Igor Pro (WaveMetrics, Lake Oswego, OR). The external solution contained 145 mM NaCl, 2.5 mM KCl, 2 mM CaCl_2_, 1 mM MgCl_2_, 10 mM N-2-hydroxyethylpiperazine-N’-2-ethanesulfonic acid (HEPES), and 11 mM glucose, adjusted to pH 7.4 with NaOH. Cells cultured on poly-L-lysine-coated glass coverslips were visually identified using an inverted microscope. Patch electrodes had tip resistances of 2–5 MΩ when filled with the internal solution (130 mM CsCl, 10 mM NaCl, 10 mM HEPES, 5 mM egtazic acid [EGTA], 4 mM MgATP, and 0.3 mM Li-GTP, adjusted to pH 7.25 with CsOH). The holding potential was set to −80 mV. After the whole-cell configuration was established, series resistance was compensated. Sodium currents were elicited using square test pulses. The voltage of the test pulse in each cell was determined by the peak of current/voltage plots that were obtained using a step voltage protocol with increments of 10 mV, from −80 to +40 mV. After a stable baseline current was obtained, anti-Nav1.7 antibody or negative control antibody solution (CP147, Bio X Cell, Inc. West Lebanon, NH) was added for 10 minutes and the effects on sodium currents were evaluated. The percentage inhibition of anti-Nav1.7 antibody or negative control antibody on Nav1.7 sodium currents was then calculated.

### Isolation of DRG cells

Rats were decapitated under deep anesthesia with isoflurane and dorsal laminectomy was performed in the lumbar region. Both L4 and L5 DRG were isolated and their connective tissues were removed in oxygenated, iced, low-sodium Ringer’s solution (212.5 mM sucrose, 3 mM KCl, 1 mM NaH_2_PO_4_, 25 mM NaHCO_3_, 11 mM D-glucose, and 5 mM MgCl_2_). The isolated DRG cells were then incubated in low-sodium Ringer’s solution containing 1–2 mg/mL collagenase for 1–2 hours at 37°C. Next, trituration was gently applied to dissociate the DRG cells in culture medium (Dulbecco’s Modified Eagle Medium with 10% fetal bovine serum, 20 mM HEPES, 1% penicillin-streptomycin solution, and 4 mM L-glutamine). After centrifugation at 1000 rpm for 5 min, the supernatant was removed and the pellet containing the DRG cells and surrounding tissue were resuspended in 10 mL of culture medium. After removing the surrounding tissue using a cell strainer and a second centrifugation at 1000 rpm for 5 minutes, the DRG cells were resuspended in 2 mL of culture medium. Next, the DRG cells were placed onto poly-L-lysine-coated glass coverslips and incubated for at least 2 hours.

### Membrane potential recording

Whole-cell patch-clamp recording was performed on visually identified DRG neurons using a Patch Clamp Amplifier (HEKA, Lambrecht, Germany). The external solution for the recording contained 145 mM NaCl, 2.5 mM KCl, 2 mM CaCl_2_, 1 mM MgCl_2_, 10 mM HEPES, and 11 mM D-glucose, adjusted to pH 7.4 with NaOH. Patch electrodes had tip resistances of 2–8 MΩ when filled with the internal solution (135 mM K gluconate, 5 mM KCl, 10 mM HEPES, 1.1 mM EGTA, 2 mM MgCl_2_, 3 mM MgATP, and 0.3 mM Li-GTP, adjusted to pH 7.2 with KOH). In the whole-cell configuration, resting membrane potentials or current-induced action potentials were measured in the current clamp mode and digitized for computer analysis using Chart software (ADInstruments, Colorado Springs, CO). Only neurons that had stable resting membrane potentials of at least −40 mV and multiple current-induced action potentials that overshot 0 mV were used for further analysis. Cells were treated with anti-Nav1.7 antibody or negative control antibody solution for 10 minutes, and the effects on the frequency of current-induced action potentials (post-treatment action potentials) were evaluated. All recordings were performed at room temperature. When resting membrane potentials during the recording period were unstable, no further measurements were conducted and the experiment was excluded from the study. The frequency of post-treatment action potentials at each stimulus was calculated.

### Animals

Five-week-old male Sprague Dawley rats were obtained from CLEA Japan (Tokyo, Japan). The animals were allowed free access to chow and tap water and were housed in a temperature-controlled room maintained on a 12-hour light/dark cycle. All experiments other than the *in vivo* extracellular recordings were conducted in compliance with the Act on Welfare and Management of Animals in Japan and the Guide for the Care and Use of Laboratory Animals, and in accordance with the protocol approved by the Institutional Animal Care and Use Committee of Shionogi & Co., Ltd., which is accredited by AAALAC International (accredited unit number: 001500). The *in vivo* extracellular recordings were conducted according to the Regulations for the Care and Use of Laboratory Animals at the University of Toyama (Approval No. A2023PHA-13 and A2020PHA-12), the Fundamental Guidelines for Proper Conduct of Animal Experiments and Related Activities in Academic Research Institutions in Japan, and the Ethical Guidelines of the International Association for the Study of Pain.

### PSNL model

The establishment of the PSNL model was as previously described (Fujita et al., 2018). Briefly, rats were anesthetized under 4% isoflurane and the skin of the left thigh was shaved and incised. After dissecting the muscle, the sciatic nerve was exposed, and the dorsal half of the nerve was ligated with a tight 4–0 nylon thread suture. The contralateral right sciatic nerve was exposed but not sutured as the sham side.

### Behavioral tests

The mechanical threshold was determined by the up-down method (Chaplan et al., 1994) using vFF ranging from 0.07–26 g (North Coast Medical, Morgan Hill, CA). Rats were placed on a mesh floor covered with an inverted transparent plastic box. A vFF was applied to the central region of the plantar surface of the hindpaw and the withdrawal response was observed. The weakest stimulation that caused a withdrawal response was taken as the PWT. The percentage reversal of the PWT was calculated as follows:

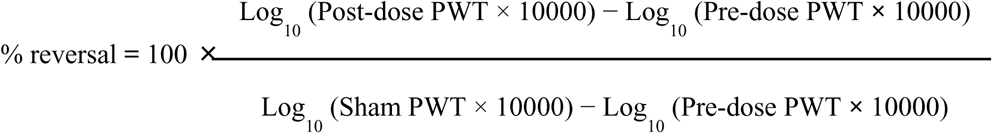

### Assessment of antibody concentrations in blood

The concentrations of antibodies in plasma/serum were measured using a Gyrolab xP workstation (Gyros Protein Technologies AB, Uppsala, Sweden). Our antibody-specific biotinylated capture peptide was added onto a Gyrolab Bioaffy 200 compact disc, which contained affinity columns preloaded with streptavidin-coated beads. Captured human IgG was then detected using an Alexa 647-anti-human IgG Fc diluted in Rexxip F buffer. The resultant fluorescent signal was recorded using the Gyrolab xP workstation.

### *In vivo* extracellular recordings from the spinal dorsal horn

*In vivo* extracellular recordings were performed as described previously (Uta et al., 2023). Rats were anesthetized with urethane (1.2–1.5 g/kg, intraperitoneal) to achieve a stable anesthetic level without the need for additional dosing. Thoracolumbar laminectomy was then performed from L4 to L5. The rats were placed in a stereotaxic apparatus and the dura mater was removed. The arachnoid membrane was then cut to create a large window on the spinal cord for the insertion of a tungsten microelectrode. The spinal cord was continuously irrigated with Krebs solution equilibrated with 95% O_2_ and 5% CO_2_ (10 –15 mL/minute) containing 117 mM NaCl, 3.6 mM KCl, 2.5 mM CaCl_2_, 1.2 mM MgCl_2_, 1.2 mM NaH_2_PO_4_, 11 mM glucose, and 25 mM NaHCO_3_ at 37°C ± 1°C. Extracellular single-unit recordings of neurons in the superficial dorsal horn (laminae I and II) were conducted at a depth ranging from 20–150 µm from the surface. Unit signals were amplified using an EX1 amplifier (Dagan Corporation, Minneapolis, MN). The recorded data were digitized using a Digidata 1400A analog-to-digital converter (Molecular Devices), stored on a personal computer using Clampex software (version 10.2; Molecular Devices), and analyzed using Clampfit software (version 10.2; Molecular Devices). To determine the specific area on the hindpaw at which a mechanical stimulus elicited a neural response, a vFF was used. A series of vFF (1.4, 4.0, 8.0, 26.0, and 60.0 g, North Coast Medical) were applied to examine the firing rates of neurons in the superficial spinal dorsal horn. Mechanical stimulation was applied for 10 s at the point of maximum response within each receptive field on the hindlimb. The percentage reversal was calculated by setting the mean value of the vehicle-treated sham group as 100% and that of the vehicle-treated PSNL group as 0%.

### Motor function assessment (rotarod test)

Sedation and motor function were tested using an accelerating rotating rod (Stoelting, Inc., Wood Dale, IL) as described previously (Khan et al., 2018; Yokoyama et al., 2007). Seven-week-old male Sprague Dawley rats were trained to stay on the rotarod at a speed of 8 rpm. On the experimental day, the rats were tested before drug administration (pre-dose) for 300 s on an accelerating rotarod at a speed that gradually increased from 4 rpm to 44 rpm. The time at which the rats fell off the rotarod was noted. The rats were then divided into four treatment groups: intravenous vehicle, intravenous anti-Nav1.7 antibody, oral vehicle, and oral pregabalin. The pregabalin (30 mg/kg) was used as a positive control as previously reported (Khan et al., 2018; Yokoyama et al., 2007). After drug administration (post-dose), the rats were again tested for 300 s on an accelerating rotarod that gradually increased from 4 rpm to 44 rpm. Anti-Nav1.7 antibody or vehicle was injected intravenously at dose of 15 mg/kg (clone1) or 10 mg/kg (S-151128), and the rotarod test was performed 5–8 hours after administration. Pregabalin or vehicle was treated orally at doses of 30 mg/kg and the rotarod test was performed 3 hours after administration.

### Immunohistochemistry

PSNL model rats under anesthesia with isoflurane were perfused transcardially with cold PBS and then formalin. Next, L4–L5 DRG were removed, post-fixed in formalin, and cryoprotected with 30% sucrose at 4°C. The DRG were then frozen in Tissue-Tek optimal cutting temperature compound (Sakura Finetech, Tokyo, Japan) and sections (10 µm) were cut using a cryostat (CM1850; Leica, Nussloch, Germany) and placed onto glass slides. For immunohistochemistry, the sections were incubated with anti-pERK antibody (1:200, #4370S, Cell Signaling Technologies, Danvers, MA) diluted in blocking solution (10% normal goat serum and 2% bovine serum albumin in PBS with Tween 20) overnight at room temperature. After being washed with PBS, the sections were incubated with anti-rabbit Alexa 488-conjugated fluorescent secondary antibody (1:500, Thermo Fisher Scientific) for 1 hour at room temperature. Sections were then mounted with VECTASHIELD (Vector Laboratories) and coverslipped. Immunostaining was visualized using a BZ-X710 microscope (KEYENCE), and pERK-positive cells in the DRG were counted using ImageJ software (National Institutes of Health, Bethesda, MD). Data were obtained for at least two sections per rat.

### Statistical analysis

All data are presented as the mean ± standard error of the mean (SEM). Significance was determined using Student’s *t*-test for non-paired samples. For multiple comparisons, analysis of variance (ANOVA) was performed followed by *post hoc* Dunnett’s test or Holm–Šidák test. GraphPad Prism (Ver. 6.07, GraphPad, Boston, MA) was used to perform statistical analyses. A value of *p* < 0.05 was considered significant.

## Acknowledgments

We thank Bronwen Gardner, PhD, from Edanz (https://jp.edanz.com/ac) for editing a draft of this manuscript.

## Author contributions

Conceptualization: S.Y., K.Y., and E.K.

Methodology: D.U., T.Y., and S.Y.

Investigation: D.U., T.Y., S.Y., K.T., T.I., S.F., D.N., and S. K.

Visualization: S.Y. and D.U.

Funding acquisition: T.T., M.Y., K.O, and E.K

Project administration: S.Y. and E.K

Supervision: E.K

Writing – original draft: S.Y. D.U.

Writing – review & editing: All authors

## The Paper Explained

### Problem

Neuropathic pain, a chronic pain condition caused by nerve injury or disease, remains difficult to treat effectively due to the limitations of existing analgesics. The Nav1.7 sodium channel is essential for pain perception, as individuals lacking Nav1.7 do not experience pain. However, the development of selective Nav1.7 inhibitors has been hindered by the high sequence similarity among sodium channel subtypes, posing a significant challenge for achieving target specificity.

### Result

To overcome this limitation, monoclonal antibodies selectively targeting Nav1.7 were developed and subsequently humanized for potential clinical application. These humanized antibodies displayed high binding affinity and selectivity toward Nav1.7 and demonstrated functional inhibition of the channel. In a rat model of neuropathic pain induced by partial sciatic nerve ligation, systemic administration of the antibody produced a potent analgesic effect lasting at least 96 hours. Electrophysiological studies confirmed that the antibody reduced mechanically evoked and spontaneous neuronal activity and suppressed mechanical stimuli–induced ERK phosphorylation in the dorsal root ganglion. Importantly, no adverse effects on physiological pain perception or motor function were observed.

### Impact

These findings suggest that humanized anti-Nav1.7 antibodies represent a promising new therapeutic approach for chronic pain. The candidate antibody S-151128 is currently undergoing clinical evaluation, highlighting its potential as a first-in-class analgesic targeting Nav1.7.

## Supplementary Figure

**Fig. S1.**
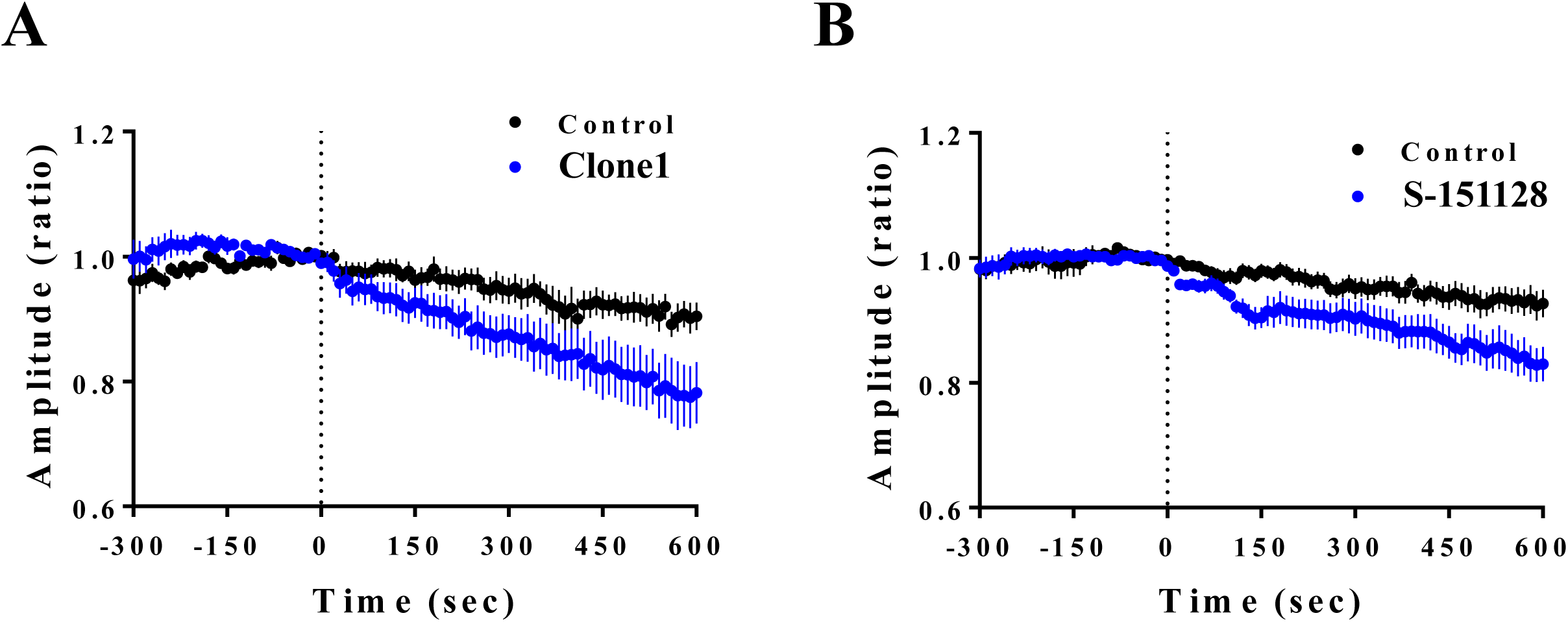
Time course of functional inhibition in rat Nav1.7. Whole cell patch clamp recordings were conducted on HEK cells expressing rat Nav1.7 to assess the inhibitory effects of antibodies at a concentration of 100 µg/mL on the sodium current. The data are presented as the mean ± SEM (n = 10 to 12). The time-course of peak currents during the experiments is shown in A and B, with the dotted lines indicating the start of antibody perfusion.

